# Population Size Mediates the Effects of Mating Systems on Evolutionary Rescue Under Worsening Environments

**DOI:** 10.64898/2026.03.30.715329

**Authors:** Neelam Porwal, Jonathan M. Parrett, Franky Rogers, Jacek Radwan, Robert J. Knell

## Abstract

Animal mating systems are hugely diverse, ranging from species where mating is essentially random to those exhibiting complex systems of mate choice by one or both sexes, with some species mating monogamously and others showing various degrees of promiscuity. There is now good evidence that if male signal traits are correlated with fitness, then polygynous female choice systems can show improved adaptation and persistence, but there has been little exploration of the ways that other types of mating systems modulate adaptation and evolutionary rescue. To address this, we developed an individual-based model that allows random mating, female-only choice, and mutual mate choice to be compared within both monogamous and polygynous frameworks and used it to explore how mating systems influence adaptive response, loss of heterozygosity, and extinction risk under changing environmental conditions. We find that mating system interacts with population size in determining extinction risk: because mate choice under polygyny lowers effective population size, accelerating the loss of heterozygosity, small polygynous populations with either mutual or female-only mate choice face higher extinction risks than randomly mating populations. In larger populations where inbreeding and genetic drift are less important, female-choice and mutual-choice polygynous systems show the greatest resilience to environmental change by allowing better-adapted males to dominate reproduction. Random mating populations show the lowest resilience to environmental change when populations are large and mutual-choice monogamous systems have intermediate resilience. Among polygynous systems, female-only choice leads to slower loss of heterozygosity and facilitates population resilience better than mutual mate choice. These findings demonstrate that mating systems can critically shape a population’s ability to adapt to environmental change and alter extinction risks, emphasizing the need to consider mating systems in designing effective conservation strategies.

## Introduction

Rapid environmental change, driven by climate change, habitat destruction, and pollution, is accelerating biodiversity loss and increasing extinction risks across taxa (1–6). Habitat fragmentation, a pervasive consequence of anthropogenic disturbance, reduces population sizes, limits gene flow, thus diminishing effective population size, leading to increased inbreeding, loss of adaptive potential, and dominance of drift over selection (7,8). Small populations are especially vulnerable to the extinction vortex, a self-reinforcing process in which demographic, environmental, and genetic factors interact to drive populations toward extinction. Understanding the mechanisms that either exacerbate or mitigate this vortex remains a central challenge in evolutionary ecology (9).

Answers to questions about population persistence or extinction in the face of environmental change have traditionally looked at ecological and demographic factors. Recently, however, theoretical and empirical studies have demonstrated an important role for mating systems in determining population viability: specifically, studies comparing random/ enforced monogamous mating versus systems in which female choice or inter-male competition can operate have found considerable differences in persistence and evolutionary rescue (10–14). Theory indicates that mate choice in a polygynous system can have beneficial effects on population persistence by favouring individuals with traits linked to higher condition or environmental match, promoting the spread of beneficial alleles or removing deleterious mutations (15,16), although the outcome may be modulated by demographic feedbacks (13) or sexual conflict (17). A meta-analysis of experimental evolution studies shows that strong sexual selection on males can improve population fitness and accelerate adaptation (18). However, comparative studies using different proxies for sexual selection strength yield inconsistent results: some report reduced population persistence under stronger sexual selection (19,20), whereas others find the opposite pattern in systems experiencing anthropogenic disturbance (21,22), and several studies detect no clear relationship (23,24). These discrepancies may reflect variation in population size, the type and duration of environmental stressors, and whether studies involve wild versus laboratory populations (25–32), but possibly could be modulated by differences in mating systems between the species analysed.

Mating systems are extraordinarily variable across the animal kingdom, with varying degrees of mate choice by both sexes plus a continuum from monogamous mating to fully polygamous systems (33–35). Monogamy tends to equalize male contributions to the breeding pool, thereby increasing effective population size and buffering populations against inbreeding and drift. In contrast, polygyny—characterized by strong reproductive monopolization by a few males and elevated male mortality—increases variance in reproductive success through both sex-ratio bias and overall demographic variance (36,37), reducing effective population size (38,39) and potentially accelerating genetic erosion (40). Mating systems can also, however, modulate the beneficial effects of sexual selection. For example, mate choice in monogamous systems compared to polygamous systems increases the reproductive contribution of less attractive, lower quality males within the mating pool (41), possibly limiting such benefits. Alternatively, mutual mate choice in monogamous systems could lead to assortative mating (42–46) with well-adapted individuals mating with each other and producing well-adapted, vigorous offspring, while poorly adapted individuals or those carrying a large mutational load would mate with each other, producing maladapted or otherwise unhealthy offspring, potentially accelerating the removal of these maladaptive alleles.

Theoretical models of sexual selection under environmental change have demonstrated their potential to increase population resilience under environmental change (13,47,48). Previous studies have also looked at the effects of demographic stochasticity on extinction risks in monogamous and polygamous mating systems (37,41,49–52). However, how monogamous and polygynous systems could alter population resilience and adaptation, while taking into consideration the direction of mate choice is not clear under environmental change. Furthermore, most models have not explicitly incorporated genetic effects such as the loss of heterozygosity in small populations, a key process involved in generating extinction vortices (53,54). Fromhage et al. (55) demonstrated that in finite populations, female choice polygamy may lead to increased inbreeding among descendants of the most successful mates but did not explore its consequences for population resilience. Without modelling a range of mating systems while accounting for the effects of co-ancestry of genomic regions, it is difficult to predict how sexual selection acting via mate choice influences long-term population persistence, particularly under worsening environments. Our work addresses these key limitations by integrating population demography, mating system structure, and population-genetic processes within a single modelling framework. With this model, we aim to evaluate extinction risks of populations differing in carrying capacities and facing environmental change by comparing (i) random mating systems versus female mate choice and mutual mate choice systems and (ii) monogamy versus polygyny.

## Results

Using an individual-based model that tracked environmental conditions, adaptation, and heterozygosity, we compared extinction risk across a range of mating systems. The environment was represented by a single environmental value (*e*), corresponding to any abiotic factor (e.g., temperature or salinity, etc.) that could remain relatively stable or vary through time. Each individual possessed a quantitative phenotype that determined its adaptation to the environment. Maladaptation was quantified as the environmental mismatch (*M*), defined as the absolute difference between an individual’s phenotype and the current environmental value. A separate finite-locus component of the model tracked heterozygosity (*H*).

Using parameter values listed in Table 1, we evaluated each mating system under varying rates of environmental changes, population sizes, costs of homozygosity, and phenotypic mismatch. Each simulation generated two primary outputs: (i) population size dynamics, quantified as the numbers of males, females, immatures, total and effective population sizes through time, and (ii) population qualities with respect to the environment, including temporal changes in *e*, mean population phenotype, *H*, and *M*.

**Table 1:** Parameter values assigned for different variables for different mating systems.

| **Choice System** | **Female**<br>**Preference**<br>**for Males**<br>***(β<sub>f</sub>)*** | **Male**<br>**Preference**<br>**for Females**<br>***(β<sub>m</sub>)*** | **Mating**<br>**System** | **Parental**<br>**Care** | **Female**<br>**Signal**<br>**Trait**<br>**Cost** | **Male**<br>**Signal**<br>**Trait**<br>**Cost** |
| --- | --- | --- | --- | --- | --- | --- |
| **Random**<br>**Monogamy** | 0 | 0 | Monogamy | FALSE | 0 | 0 |
| **Mutual Choice**<br>**Monogamy** | 5 | 5 | Monogamy | TRUE | 0.25 | 0.25 |
| **Random Polygyny** | 0 | 0 | Polygyny | FALSE | 0 | 0 |
| **Female Choice**<br>**Polygyny** | 5 | 0 | Polygyny | FALSE | 0 | 0.25 |
| **Polygyny with**<br>**Mutual Choice** | 5 | 5 | Polygyny | FALSE | 0.25 | 0.25 |

### Stable environments

Under relatively stable environmental conditions (rate of directional change = 0, Fig 1), with only very small fluctuations in *e, H* declined over generations across all mating systems and parameter combinations. The homozygosity penalty determined how strongly individuals with reduced *H* were penalized and consequently how strongly selection acted against individuals with common ancestry. The decline in heterozygosity through time depended on the penalty and was steepest when the homozygosity penalty was set to zero. Across ranges of homozygosity penalties, polygynous systems with mate choice showed the steepest declines in *H* in line with their low *N*_*e*_ resulting from the reproductive skew among males characteristic of this system. In stable environments, extinctions occurred only at small and, much less frequently, at intermediate population sizes. Severe homozygosity penalties increased extinction risk, and systems with mate choice were most prone to faster and higher extinction rates (Fig1).

**Fig 1.**
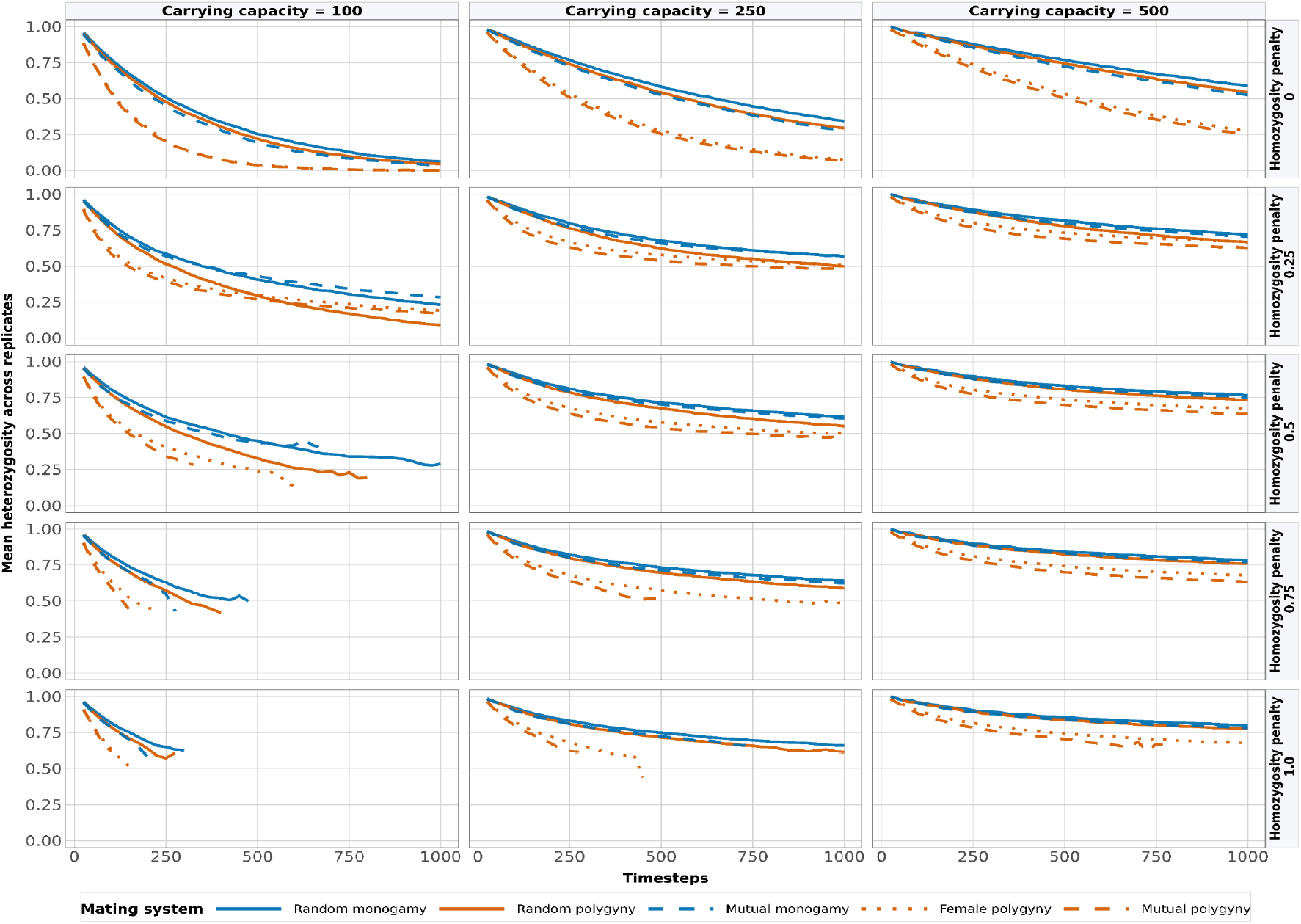
Median (population) heterozygosity averaged over 50 simulations per parameter combination. Simulations were run across a thousand timesteps, under different mating systems with no environmental change and zero mismatch penalty (*M* = 0). Vertical panels represent different carrying capacities (*K*), while horizontal panels show varying homozygosity penalties (*H*). Heterozygosity is calculated only for populations that persisted (i.e., did not go extinct) until respective timesteps. Blue lines represent monogamy and orange lines represent polygyny in mating systems. Random, mutual choice and female choice mating systems are indicated by solid, dashed and dotted lines respectively. Each parameter combination was run 50 times and the median population heterozygosity was recorded for every 25 time steps.

### Changing environments

Under changing-environment scenarios (directional change > 0), *e* was kept relatively stable with small fluctuations during the initial 50 generations, allowing populations to stabilise, and then began to change. During this initial period, the mean phenotype closely matched the environmental optimum. Once *e* began to shift directionally, phenotypes followed but lagged behind the changing optimum, in some scenarios leading to *M* increasing considerably with time. Rising *M* sometimes reduced population sizes sufficiently to cause extinction (Fig. 2, S2, S3). At *K* = 500 and a homozygosity penalty of 0.5, population sizes declined more drastically under random mating compared to mutual-choice and female-choice systems (Fig S3A), but in small populations (*K* = 100) with a homozygosity penalty of 1, populations under random monogamy persisted for a longer time (Fig S2B).

**Fig 2:**
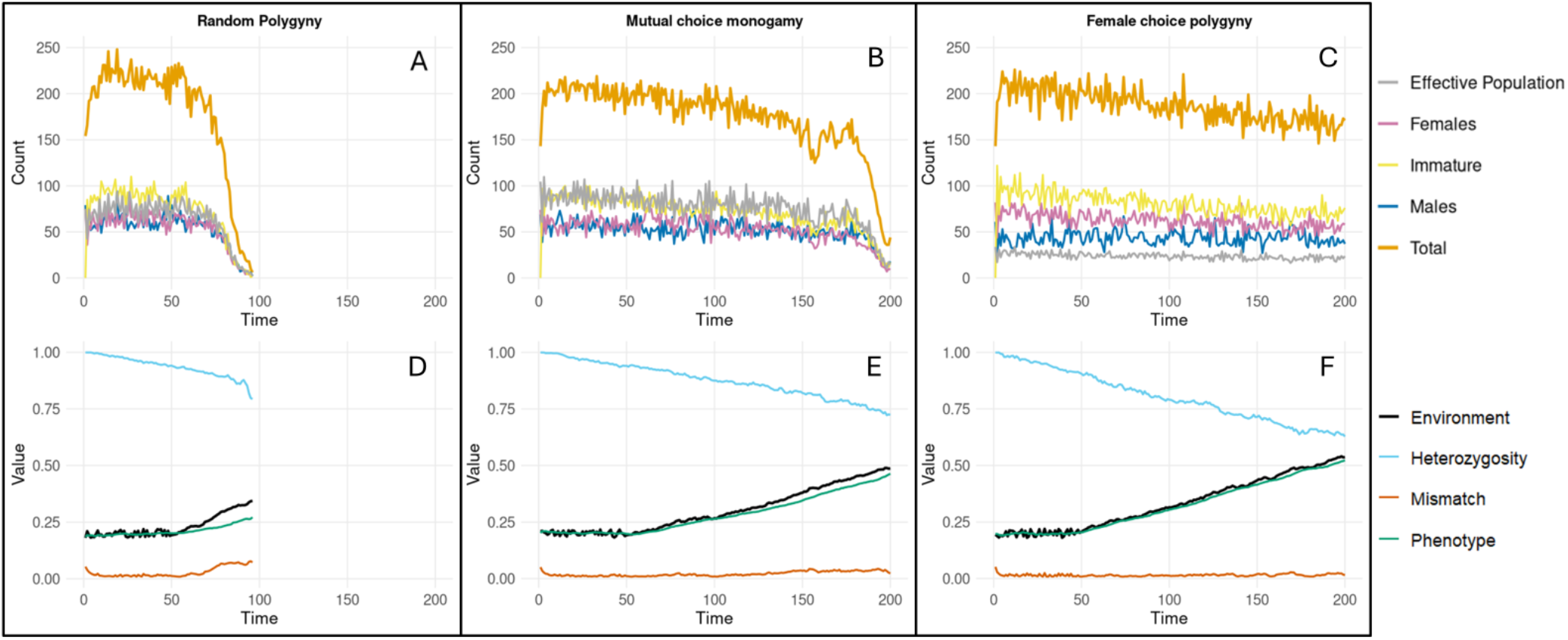
Example outputs from simulation runs for each type of mating system when directional rate = 0.002. A-C tracking the population demography. D-F tracks the environment, and the population qualities with respect to the environment. A and D, shows random mating polygyny. B and E, shows mutual choice monogamy. C and F, shows female choice polygyny. All three simulation runs were run with K = 250, time steps = 200, mismatch penalty *=* 5 and inbreeding penalty = 0.5.

The proportions of populations that went extinct and median times to extinction at different levels of *M* and homozygosity penalty are shown in Fig S4. There was a general trend of lower extinction risks in female-choice polygynous systems compared to other systems at high *K* and, at medium and low *K*, under lower homozygosity penalties. To visualise general trends, we calculated a value referred to as resilience, defined as the rate of environmental change predicted to cause population extinction 50% of the time; thus, higher values indicate a better tolerance for environmental change. Conversely, a value of zero for resilience indicates extinction occurring for that combination of parameter values even when the environment is stable.

Fig 3 shows resilience values across all combinations of *K*, mismatch penalty, homozygosity penalty, and mating system. A complex interaction between mating systems, population size, and homozygosity penalty was apparent. For all the population sizes modelled, when the penalty for homozygosity was close to zero, female- and mutual-choice polygyny systems had the highest resilience and random mating systems had the lowest, with mutual-choice monogamy being intermediate. As the penalty for homozygosity increased, however, these differences either disappeared (in large populations) or reversed (in small populations), with mutual-choice polygyny showing the lowest resilience in small-to-medium populations, female-choice polygyny and mutual choice monogamy converging on similar, low resilience values and the two random mating options having the highest resilience. This interaction between population size and the cost of homozygosity was more pronounced when the effect of phenotypic distance from the environmental optimum (the mismatch penalty) was higher.

**Fig 3:**
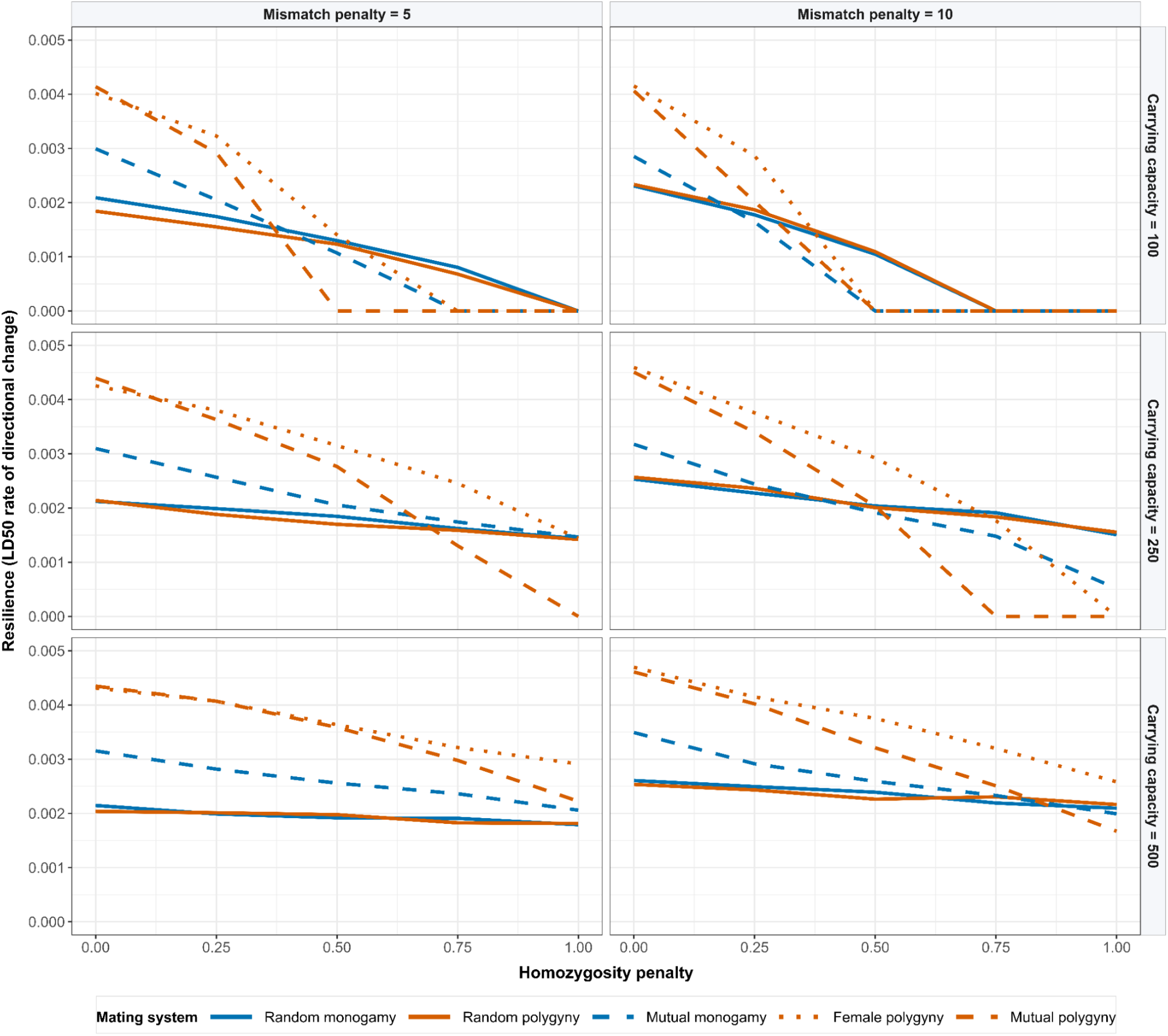
Rate of environmental change, predicted to result in 50% extinction expressed as resilience, across parameter combinations and mating systems. Homozygosity penalties (*H*) are plotted on the x-axis. Vertical panels represent different mismatch penalties (*M*), while horizontal panels show varying carrying capacities (*K*). Blue lines represent monogamy and orange lines represent polygyny in mating systems. Random, mutual choice and female choice mating systems are indicated by solid, dashed and dotted lines respectively. Each parameter combination was run 50 times.

## Discussion

The capacity of a population to adapt to environmental change and to maintain heterozygosity is a crucial component of the persistence of small populations, but the role of the diversity of mating systems found across the animal kingdom in shaping this capacity remains insufficiently understood. To address this, we examined how population size, costs of heterozygosity loss, and the nature of the mating system interact under environmental change to determine persistence using an individual-based simulation model. In broad terms, our model shows that mate-choice systems enhance resilience in large populations but may reduce it in small populations due to accelerated loss of heterozygosity. Our results further reveal that within mate-choice-based systems, the monogamy-polygamy axis is a key determinant: populations under polygynous systems exhibited consistently higher population survival rates than monogamous systems, except when the cost of homozygosity was high, when mutual choice polygyny had lower resilience. Within polygyny, female choice conferred a higher resilience compared to mutual choice across all tested scenarios.

Previous theoretical works have largely compared female-choice polygyny with systems lacking mate choice but have not examined a broader array of mating systems or, critically, the consequences of mate choice for heterozygosity loss in small populations. Our simulations show that both these factors may have key consequences for population resilience. Polygynous systems exhibited the most pronounced declines in heterozygosity over successive generations (Fig 1, S2, S3). As a consequence of the reduction in *N*_*e*_ that arises from reproductive skew in these systems. In contrast, mutual choice monogamous systems show comparatively larger *N*_*e*_ and slower declines in heterozygosity, despite both sexes experiencing increased mortality due to costs of signal trait expression at the same level as in mate-choice based polygyny systems (Fig 1, S2, S3). This pattern suggests that the more substantial reduction in heterozygosity observed in polygynous populations is driven primarily by reproductive skew among males. At low population sizes, this accelerated decline in heterozygosity, and consequent exposure of recessive genetic load, may have particularly severe consequences, leading to extinction in small populations even in the absence of additional stress, as seen in our model, even in the absence of environmental change over 1000 generations.

Mate-choice based polygyny allows a rapid adaptive response because the best adapted males achieve disproportionally high reproductive success (13,15). Under sustained directional environmental change, this more than offsets the negative consequences of mate-choice-based polygyny on *Ne* and heterozygosity. In large populations, therefore, or when homozygosity penalty was low, mate-choice based polygyny resulted in better resilience compared to monogamy (Fig 3). This increased rate of adaptation is, however, insufficient to buffer small populations from extinction when homozygosity penalty is high, hence the lower resilience shown by mate choice polygynous systems. These findings highlight how demographic decline, heterozygosity erosion, and reduced adaptive capacity interact to determine persistence under environmental stress. They also help explain why empirical and comparative studies report inconsistent effects of sexual selection on population viability (10–13,19,20). Our work suggests that future analyses should include mating systems, population size, and heterozygosity as potential variables accounting for these varying outcomes.

Monogamy is common in birds (56) and in many other taxa (57), and mutual mate choice likely occurs in many of these systems. An important question regarding this understudied mating system is how it affects population persistence, especially under environmental change. As mentioned in the introduction, with mutual mate choice, there is the possibility for well-adapted males and females to pair up and thereby produce extremely well-adapted “super” offspring, but whether this would increase adaptation more than in, for example, female choice polygyny is not intuitively clear. Our results show that when the cost of homozygosity is small, or populations are relatively large, mutual choice monogamy leads to resilience that is intermediate between random mating and mate-choice polygyny systems. In small populations with high homozygosity penalties, the additional mortality arising from signal-trait expression in both sexes can diminish the resilience of mutual-choice monogamous systems, making them less robust than random-mating populations where such costs do not occur. One limitation of the present model that is relevant here is that it does not include the potential for extra-pair copulations, which are common in many socially monogamous systems (58), which could alter this result; this will be the subject of a future study.

Differences in the direction and symmetry of mate choice further refine our understanding of these broader dynamics. While comparing polygynous systems, in systems with female-only choice, signalling costs fall solely on males, increasing male mortality without directly affecting female survival. In mutual-choice systems, signalling costs occur in both sexes (59–61), producing more balanced sex ratios and reduced reproductive skew (62,63). However, costs incurred by females result in higher female mortality under mutual mate choice compared to female-only choice (Fig. 2, S2, S3). Since reproductive output is limited by the number of surviving females, mutual-choice systems consequently may generate fewer offspring than female-choice systems, increasing the risk of populations not being able to maintain the size admitted by carrying capacity.

Previous models investigating extinction risk in small populations under stable conditions have likewise shown that elevated female mortality reduces resilience and increases vulnerability to demographic stochasticity (49–51). These outcomes due to elevated female mortality are further highlighted in the recent empirical work under environmental change (64). Our model extends these findings by incorporating heterozygosity dynamics, which earlier behavioural models did not include. We note that a similar level of female mortality to that in mutual-choice polygamy occurs in mutual-choice monogamy. The reasons why these two systems nevertheless differ in resilience further attest to the role of reproductive bias causing these differences, as discussed above.

Ultimately, whether a mating system enhances resilience or increases extinction risk depends on how it interacts with population size and genetic processes. Our findings show that mating systems can shape demographic and evolutionary trajectories in contrasting ways, sometimes accelerating adaptation, and other times amplifying vulnerability through reduced N_*e*_, increased heterozygosity loss, or sex-specific mortality. Recognizing this duality is essential for forecasting population persistence under environmental change. Our model does not incorporate sexual conflict, which may alter population responses (17,64–66). Future extensions should include additional mating systems such as polyandry and social monogamy. Integrating mating-system diversity, reproductive skew, sex-specific costs, and mating opportunity structure will improve predictions of species resilience and inform conservation strategies in rapidly changing environments.

## Methods

We developed an individual-based simulation model to explore the dynamics of mate choice strategies, including symmetric (mutual mate choice), asymmetric (female-only choice), and random mating, with monogamy and polygyny under varying environmental conditions. The model simulates a spatially homogeneous population of individuals defined by their genetic architecture and sexual traits, evolving over discrete time steps, following the ODD protocol for individual-based models (67). Implemented in R (version 4.5.2)(68), the model operates across three hierarchical levels (individual, population, and environment), each characterized by state variables that interact dynamically. Variables and formulas used are additionally described in tables S1 and S2, and the schematic flow of the model is described in Figure S1.

The model is coded in a single function, consistent with R-based simulation guidelines (69). The population starts with an initial number of individuals equal to the carrying capacity K, with probabilistically likely balanced ages uniformly distributed between 1 and 10. Each individual is defined by state variables: survival status (alive or dead), sex, age, genotype, phenotype, heterozygosity, environmental mismatch, and a sex-specific signal trait. In order to model both adaptation and changes in heterozygosity within the manageable computational power required, we use a mixed framework to model the genetic architecture of the system. Each individual thus has two heritable components to its biology: the single value that determines adaptation to the changing environment and an array of markers that trace co-ancestry of unlinked genomic regions that allows us to model changes in heterozygosity.

Traits such as thermal tolerance are often highly polygenic: as an example, Williams-Simon et al. (70) compared RNA-seq data for high and low thermal tolerance lines of *Drosophila melanogaster* and found differential expression at 2,642 loci. A finite-locus model with a realistic number of loci in an individual-based framework would be computationally challenging, hence our adoption of this mixed approach. To model adaptation, we use an approach based on Fisher’s infinitesimal model. Phenotype is a continuous trait drawn from a normal distribution (mean = 0.2, sd = 0.1) at initialization, representing a trait under selection, with the optimum value depending on the environment (see below). The offspring phenotype was inherited as the average of parental phenotypes plus a normal mutation (sd = 0.01).

To model the effects of loss of heterozygosity, following Fromhage et al. (55), we included a component consisting of 50 diploid unlinked loci, each with two alleles (unique integers from 1 to 1000), initialized randomly. Offspring genotypes were generated following Mendelian inheritance principles, in which one allele is randomly inherited from either of the strands from each parent at every locus (71). This was used to calculate heterozygosity (*H*) as the proportion of loci where the two allelic strands differ. 1-*H* thus measured co-ancestry across unlinked genomic regions, and thus the probability of deleterious recessives in these regions being homozygous and thus exposed to selection, but the loci themselves did not contribute to the individual’s phenotype when considering environmental adaptation. We assume the populations we model to be a result of a fragmentation with initial high population sizes under stable environments evolving for some time. We therefore initiate our populations as completely outbred ones with a starting heterozygosity of ~ 1. We aim to track the population right after fragmentation has occurred.

The environment is represented by a single variable, *e*, initialized at 0.2. Environmental *M* is the absolute difference between an individual’s phenotype and *e* (7). Upon reaching sexual maturity (age ≥ I = 2), individuals develop a sex-specific signal trait, calculated as:

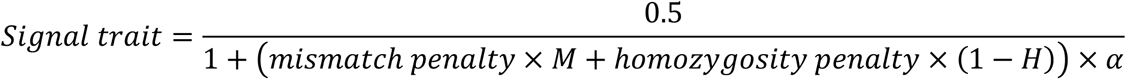

where the mismatch penalty and homozygosity penalty are variable parameters scaling environmental mismatch and heterozygosity effects, and α (condition dependence, α_m_ for males, α_f_ for females, both set to 2) determines the trait’s sensitivity to these factors. The homozygosity penalty in our model can be interpreted as the effect which exposure of deleterious recessives to selection has on fitness, i.e. the strength of inbreeding depression (72).

At each time step, the environmental variable *e* changes according to the rate of directional change. For directional rate set to zero *e* fluctuates randomly (uniform distribution, range ±0.02); otherwise, it follows the same random fluctuation for the first 50 time steps and then increases with a mean directional rate plus random variation (normal distribution, standard deviation ±0.003).

Extinction occurs if the population size reaches 0 or no individuals of one sex remain, at which point the simulation terminates, and the extinction time is recorded (7).

Each time step, individuals’ ages increase by one, and mismatch and signal traits are recalculated based on the current *e*. Survival probability is determined as:

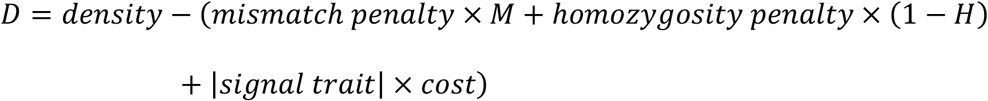

where density = *K* / population size, mismatch penalty and inbreeding penalty are variable, and cost (*cost*.*m* for males, *cost*.*f* for females) represents the survival cost of expressing the signal trait. Immature individuals (age < *I*) are exempt from signal trait costs. Individuals die if a random draw (0 to 1) is less than or equal to their death probability. Dead individuals are removed, and the population is checked for extinction each time step.

Mature individuals (age ≥ *I*) form mating pairs based on the mating system (mating system, either monogamy or polygyny) and signal trait preferences (*β*_*m*_ for males, *β*_*f*_ for females). Adjusting the values of *β* allows us to determine the mating system. For example, if *β*_*m*_ *= 0* and *β*_*f*_ *= 5*, the system is one with female choice but no male choice; if both *β*_*m*_ and *β*_*f*_ are equal to zero, the mating is random, and if both are 5, there is a degree of mutual choice. A cost for the signal trait expression of 0.25 was borne by the sex opposite to the one exerting the preference; otherwise, the cost was set to zero, resulting in the signal trait not having any effect even if it was calculated.

Females are assigned to mating groups (maximum number of females in a group = 10), and males are randomly distributed across these groups. For female choice systems (*β*_*f*_ = 5, *β*_*m*_ = 0), males assigned to each group are ranked, highest to lowest, by the degree of expression of the signal trait, and for mutual choice systems (*β*_*f*_ =50, *β*_*m*_ = 0), both males and females are ranked (73). Females select mates with probabilities:

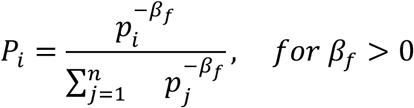

or uniformly (1 / group size) if *β*_*f*_ = 0, *P*_*i*_ is the probability that the female chooses male I, and *p*_*i*_ is the male’s rank. Males follow a similar process if *β*_*m*_ > 0.

In the monogamy treatment, group sizes were equalized by removing the more abundant sex with the lowest signal trait values in the group to ensure one-to-one pairing. In the polygyny with female choice, males were allowed to mate with multiple females, whereas each female mated only once in all mating systems. This model does not include polyandry or female-choice monogamy, because our model structure would have produced outputs equivalent to polygyny (for polyandry) or random monogamy with costly signalling (for female-choice monogamy), the latter being unlikely to represent an evolutionarily stable strategy.

Fecundity is calculated as:

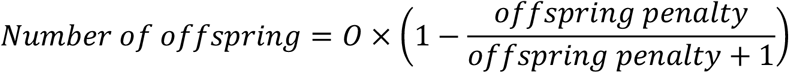

where *O* = 5 is the maximum offspring. In monogamous systems with mutual mate choice, offspring fecundity and survival depend on the contributions of both parents (74), as mutual choice arises from shared parental investment in offspring (75–80). Accordingly, we modelled parental care as an offspring penalty determined by the mismatch and heterozygosity of both parents in mutual-choice monogamous systems. In contrast, in all other mating systems, the penalty depended only on maternal traits.

Biparental care:

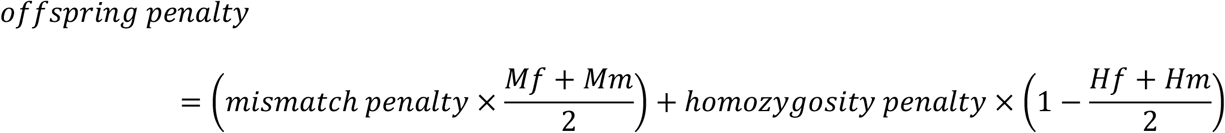

Uniparental care:

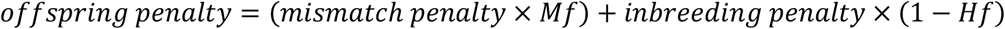

Where *Mf* is the mother’s mismatch, *Mm* is the father’s mismatch, *Hf* is the mother’s heterozygosity, and *Hm* is the father’s heterozygosity.

The actual number of newborns is then determined via a binomial process with birth probability b = 0.5 (to keep the number of offspring in check according to *K*). Offspring are added to the population, and the effective population size is computed based on the variance in reproductive success among males and females (adults).

All simulations were run on the University of Hull “Viper” high-performance cluster using the R packages foreach (81) and doparallel (82). The final simulation runs were replicated 50 times for each parameter combination for all mating systems. The parameters for mating systems were set according to Table 1. The model was run for all mating systems (Table 1) with a range of carrying capacities, rates of directional change, mismatch, and inbreeding penalties (as described in Table S1).

## Acknowledgements and Funding sources

We would like to thank Mateusz Konczal for his insights on the population genetics part of the code. The project was funded by a grant from NCN UMO-2020/39/B/NZ8/00152/4 to J. Radwan, R.J. Knell and T.C. Cameron.

## References

1. Parmesan C, Yohe G. A globally coherent fingerprint of climate change impacts across natural systems. nature. 2003;421(6918):37–42.

2. Thomas CD, Cameron A, Green RE, Bakkenes M, Beaumont LJ, Collingham YC, et al. Extinction risk from climate change. Nature. 2004;427(6970):145–8.

3. Bellard C, Bertelsmeier C, Leadley P, Thuiller W, Courchamp F. Impacts of climate change on the future of biodiversity. Ecol Lett. 2012 Apr;15(4):365–77. doi:10.1111/j.1461-0248.2011.01736.x

4. Pimm SL, Jenkins CN, Abell R, Brooks TM, Gittleman JL, Joppa LN, et al. The biodiversity of species and their rates of extinction, distribution, and protection. Science. 2014 May 30;344(6187):1246752. doi:10.1126/science.1246752

5. Ceballos G, Ehrlich PR, Barnosky AD, García A, Pringle RM, Palmer TM. Accelerated modern human–induced species losses: Entering the sixth mass extinction. Sci Adv. 2015 Jun 5;1(5):e1400253. doi:10.1126/sciadv.1400253

6. Urban MC. Accelerating extinction risk from climate change. Science. 2015 May;348(6234):571–3. doi:10.1126/science.aaa4984

7. Brook BW, Sodhi NS, Bradshaw CJ. Synergies among extinction drivers under global change. Trends Ecol Evol. 2008;23(8):453–60.

8. Spielman D, Brook BW, Frankham R. Most species are not driven to extinction before genetic factors impact them. Proc Natl Acad Sci. 2004 Oct 19;101(42):15261–4. doi:10.1073/pnas.0403809101

9. Hoffmann AA, Sgrò CM. Climate change and evolutionary adaptation. Nature. 2011;470(7335):479–85.

10. Lumley AJ, Michalczyk Ł, Kitson JJ, Spurgin LG, Morrison CA, Godwin JL, et al. Sexual selection protects against extinction. Nature. 2015;522(7557):470–3.

11. Martinossi-Allibert I, Savković U, \DJor\djević M, Arnqvist G, Stojković B, Berger D. The consequences of sexual selection in well-adapted and maladapted populations of bean beetles. Evolution. 2018;72(3):518–30.

12. Berger D, Liljestrand-Rönn J. Environmental complexity mitigates the demographic impact of sexual selection. Ecol Lett. 2024 Jan;27(1):e14355. doi:10.1111/ele.14355

13. Martínez-Ruiz C, Knell RJ. Sexual selection can both increase and decrease extinction probability: reconciling demographic and evolutionary factors. Bassar R, editor. J Anim Ecol. 2017 Jan;86(1):117–27. doi:10.1111/1365-2656.12601

14. Godwin JL, Lumley AJ, Michalczyk Ł, Martin OY, Gage MJG. Mating patterns influence vulnerability to the extinction vortex. Glob Change Biol. 2020;26(8):4226–39. doi:10.1111/gcb.15186

15. Lorch PD, Proulx S, Rowe L, Day T. Condition-dependent sexual selection can accelerate adaptation. Evol Ecol Res. 2003;5(6):867–81.

16. Whitlock MC, Agrawal AF. Purging the genome with sexual selection: reducing mutation load through selection on males. Evolution. 2009;63(3):569–82.

17. Flintham EO, Savolainen V, Mullon C. Male harm offsets the demographic benefits of good genes. Proc Natl Acad Sci. 2023 Mar 7;120(10):e2211668120. doi:10.1073/pnas.2211668120

18. Cally JG, Stuart-Fox D, Holman L. Meta-analytic evidence that sexual selection improves population fitness. Nat Commun. 2019;10(1):2017.

19. Bro-Jørgensen J. Will their armaments be their downfall? Large horn size increases extinction risk in bovids: Large horn size increases extinction risk in bovids. Anim Conserv. 2014 Feb;17(1):80–7. doi:10.1111/acv.12062

20. Martins MJF, Puckett TM, Lockwood R, Swaddle JP, Hunt G. High male sexual investment as a driver of extinction in fossil ostracods. Nature. 2018;556(7701):366–9.

21. Moore MP, Leith NT, Fowler-Finn KD, Medley KA. Human-modified habitats imperil ornamented dragonflies less than their non-ornamented counterparts at local, regional, and continental scales. Ecol Lett. 2024 Jun;27(6):e14455. doi:10.1111/ele.14455

22. Parrett JM, Mann DJ, Chung AYC, Slade EM, Knell RJ. Sexual selection predicts the persistence of populations within altered environments. Grether G, editor. Ecol Lett. 2019 Oct;22(10):1629–37. doi:10.1111/ele.13358

23. Morrow EH, Fricke C. Sexual selection and the risk of extinction in mammals. Proc R Soc Lond B Biol Sci. 2004;271(1555):2395–401.

24. Morrow EH, Pitcher TE. Sexual selection and the risk of extinction in birds. Proc R Soc Lond B Biol Sci. 2003;270(1526):1793–9.

25. Bürger R, Lynch M. EVOLUTION AND EXTINCTION IN A CHANGING ENVIRONMENT: A QUANTITATIVE-GENETIC ANALYSIS. Evolution. 1995 Feb;49(1):151–63. doi:10.1111/j.1558-5646.1995.tb05967.x

26. Kirkpatrick M, Barton NH. Evolution of a Species’ Range. Am Nat. 1997 Jul;150(1):1–23. doi:10.1086/286054

27. Boulding EG, Hay T. Genetic and demographic parameters determining population persistence after a discrete change in the environment. Heredity. 2001;86(3):313–24.

28. Candolin U, Heuschele J. Is sexual selection beneficial during adaptation to environmental change? Trends Ecol Evol. 2008;23(8):446–52.

29. Holman L, Kokko H. The consequences of polyandry for population viability, extinction risk and conservation. Philos Trans R Soc B Biol Sci. 2013 368(1613).

30. Doherty PF, Sorci G, Royle JA, Hines JE, Nichols JD, Boulinier T. Sexual selection affects local extinction and turnover in bird communities. Proc Natl Acad Sci. 2003 May 13;100(10):5858–62. doi:10.1073/pnas.0836953100

31. Sorci G, Møller AP, Clobert J. Plumage dichromatism of birds predicts introduction success in New Zealand. J Anim Ecol. 1998 Mar;67(2):263–9. doi:10.1046/j.1365-2656.1998.00199.x

32. McLain DK, Moulton MP, Redfearn TP. Sexual selection and the risk of extinction of introduced birds on oceanic islands. Oikos. 1995;27–34.

33. Klug H. Animal mating systems. eLS. 2011.

34. Kokko H, Klug H, Jennions MD. Mating systems. In: Shuker DM, Simmons LW, editors. The Evolution of Insect Mating Systems. Oxford: Oxford Univ Press; 2014. p. 42–60.

35. Shuster SM, Wade MJ. Mating Systems and Strategies. Princeton: Princeton Univ Press; 2019.

36. Sæther BE, Engen S. Towards a predictive conservation biology: the devil is in the behaviour. Philos Trans R Soc B Biol Sci. 2019;374(1781).

37. Sæther BE, Engen S, Lande R, Møller AP, Bensch S, Hasselquist D, et al. Time to extinction in relation to mating system and type of density regulation in populations with two sexes. J Anim Ecol. 2004;925–34.

38. Nunney L. THE INFLUENCE OF MATING SYSTEM AND OVERLAPPING GENERATIONS ON EFFECTIVE POPULATION SIZE. Evolution. 1993 Oct;47(5):1329–41. doi:10.1111/j.1558-5646.1993.tb02158.x

39. Stiver JR, Apa AD, Remington TE, Gibson RM. Polygyny and female breeding failure reduce effective population size in the lekking Gunnison sage-grouse. Biol Conserv. 2008;141(2):472–81.

40. Kokko H, Brooks R. Sexy to die for? Sexual selection and the risk of extinction. In: Annales Zoologici Fennici. JSTOR; 2003. p. 207–19.

41. Bessa-Gomes C, Legendre S, Clobert J. Allee effects, mating systems and the extinction risk in populations with two sexes. Ecol Lett. 2004 Sep;7(9):802–12. doi:10.1111/j.1461-0248.2004.00632.x

42. Baldauf SA, Kullmann H, Schroth SH, Thünken T, Bakker TC. You can’t always get what you want: size assortative mating by mutual mate choice as a resolution of sexual conflict. BMC Evol Biol. 2009;9(1):129. doi:10.1186/1471-2148-9-129

43. Beeching SC, Hopp AB. Male mate preference and size-assortative pairing in the convict cichlid. J Fish Biol. 1999 Nov;55(5):1001–8. doi:10.1111/j.1095-8649.1999.tb00735.x

44. Greenway R, Drexler S, Arias-Rodriguez L, Tobler M. Adaptive, but not condition-dependent, body shape differences contribute to assortative mating preferences during ecological speciation. Evolution. 2016 Dec;70(12):2809–22. doi:10.1111/evo.13087

45. Harari AR, Handler AM, Landolt PJ. Size-assortative mating, male choice and female choice in the curculionid beetle Diaprepes abbreviatus. Anim Behav. 1999 Dec;58(6):1191–200. doi:10.1006/anbe.1999.1257

46. McKaye KR. Mate choice and size assortative pairing by the cichlid fishes of Lake Jiloá, Nicaragua. J Fish Biol. 1986 Dec;29(sA):135–50. doi:10.1111/j.1095-8649.1986.tb05005.x

47. Tanaka Y. Sexual Selection Enhances Population Extinction in a Changing Environment. J Theor Biol. 1996 Jun 7;180(3):197–206. doi:10.1006/jtbi.1996.0096

48. Knell RJ, Parrett JM. Alternative reproductive tactics and evolutionary rescue. Evol Lett. 2024;8(4):539–49.

49. Legendre S, Clobert J, Møller AP, Sorci G. Demographic Stochasticity and Social Mating System in the Process of Extinction of Small Populations: The Case of Passerines Introduced to New Zealand. Am Nat. 1999 May;153(5):449–63. doi:10.1086/303195

50. Møller AP, Legendre S. Allee effect, sexual selection and demographic stochasticity. Oikos. 2001 Jan;92(1):27–34. doi:10.1034/j.1600-0706.2001.920104.x

51. Bessa-Gomes C, Clobert J, Legendre S, Møller AP. Modeling Mating Patterns Given Mutual Mate Choice: The Importance of Individual Mating Preferences and Mating System. J Biol Syst. 2003 Sep;11(03):205–19. doi:10.1142/S0218339003000853

52. Leach D, Shaw AK, Weiss-Lehman C. Stochasticity in social structure and mating system drive extinction risk. Ecosphere. 2020 Feb;11(2):e03038. doi:10.1002/ecs2.3038

53. Nabutanyi P, Wittmann MJ. Models for Eco-Evolutionary Extinction Vortices under Balancing Selection. Am Nat. 2021 Mar 1;197(3):336–50. doi:10.1086/712805

54. Blomqvist D, Pauliny A, Larsson M, Flodin LÅ. Trapped in the extinction vortex? Strong genetic effects in a declining vertebrate population. BMC Evol Biol. 2010;10(1):33. doi:10.1186/1471-2148-10-33

55. Fromhage L, Kokko H, Reid JM. Evolution of Mate Choice for Genome-Wide Heterozygosity. Evolution. 2009;63(3):684–94. doi:10.1111/j.1558-5646.2008.00575.x

56. Cockburn A. Prevalence of different modes of parental care in birds. Proc R Soc B Biol Sci. 2006;273(1592):1375–83.

57. Kvarnemo C. Why do some animals mate with one partner rather than many? A review of causes and consequences of monogamy. Biol Rev. 2018;93(4):1795–812. doi:10.1111/brv.12421

58. Brouwer L, Griffith SC. Extra-pair paternity in birds. Mol Ecol. 2019 Nov;28(22):4864–82. doi:10.1111/mec.15259

59. Amundsen T. Why are female birds ornamented? Trends Ecol Evol. 2000;15(4):149–55.

60. Zahavi A, Zahavi A. The Handicap Principle: A Missing Piece of Darwin’s Puzzle. Oxford: Oxford Univ Press; 1997.

61. Hooper PL, Miller GF. Mutual Mate Choice Can Drive Costly Signaling Even Under Perfect Monogamy. Adapt Behav. 2008 Feb;16(1):53–70. doi:10.1177/1059712307087283

62. Fromhage L, Jennions MD. Coevolution of parental investment and sexually selected traits drives sex-role divergence. Nat Commun. 2016;7(1):12517.

63. McDonald GC, Pizzari T. Structure of sexual networks determines the operation of sexual selection. Proc Natl Acad Sci. 2018 Jan 2;115(1). doi:10.1073/pnas.1710450115

64. Pandey N, Porwal N, Parrett JM, Radwan J, Knell RJ, Cameron TC. Sexual selection associated with an aggressive male phenotype reduces population size and hinders population recovery after heat stress. Ecology Letters. 2026 Apr;29(4):e70377.

65. Plesnar-Bielak A, Łukasiewicz A. Sexual conflict in a changing environment. Biol Rev. 2021 Oct;96(5):1854–67. doi:10.1111/brv.12728

66. Gómez-Llano M, Faria GS, García-Roa R, Noble DW, Carazo P. Male harm suppresses female fitness, affecting the dynamics of adaptation and evolutionary rescue. Evol Lett. 2024;8(1):149–60.

67. Grimm V, Berger U, DeAngelis DL, Polhill JG, Giske J, Railsback SF. The ODD protocol: a review and first update. Ecol Model. 2010;221(23):2760–8.

68. R Core Team. R: A language and environment for statistical computing. R Foundation for Statistical Computing; 2023.

69. Acerbi A, Mesoudi A, Smolla M. Individual-based models of cultural evolution: a step-by-step guide using R. Routledge; 2022.

70. Williams-Simon PA, Oster C, Moaton JA, Ghidey R, Ng’oma E, Middleton KM, et al. Naturally segregating genetic variants contribute to thermal tolerance in a Drosophila melanogaster model system. Genetics. 2024;227(1):iyae040.

71. Falconer DS. Introduction to quantitative genetics. Pearson Education India; 1996.

72. Charlesworth D, Willis JH. The genetics of inbreeding depression. Nat Rev Genet. 2009;10(11):783–96.

73. Petrie M, Tim H, Carolyn S. Peahens prefer peacocks with elaborate trains. Anim Behav. 1991;41(2):323–31.

74. Courtiol A, Etienne L, Feron R, Godelle B, Rousset F. The Evolution of Mutual Mate Choice under Direct Benefits. Am Nat. 2016 Nov;188(5):521–38. doi:10.1086/688658

75. Parker GA. Mate quality and mating decisions. Mate Choice. 1983;141:166.

76. Crowley PH, Travers SE, Linton MC, Cohn SL, Sih A, Sargent RC. Mate Density, Predation Risk, and the Seasonal Sequence of Mate Choices: A Dynamic Game. Am Nat. 1991 Apr;137(4):567–96. doi:10.1086/285184

77. Johnstone RA, Reynolds JD, Deutsch JC. Mutual Mate Choice and Sex Differences in Choosiness. Evolution. 1996;50(4):1382–91. doi:10.1111/j.1558-5646.1996.tb03912.x

78. Kokko H, Johnstone RA. Why is mutual mate choice not the norm? Operational sex ratios, sex roles and the evolution of sexually dimorphic and monomorphic signalling. Philos Trans R Soc Lond B Biol Sci. 2002;357(1419):319–30.

79. Kokko H, Jennions MD. Parental investment, sexual selection and sex ratios. J Evol Biol. 2008;21(4):919–48.

80. Owens IPF, Burke T, Thompson DB. Extraordinary sex roles in the Eurasian dotterel: female mating arenas, female-female competition, and female mate choice. Am Nat. 1994;144:76–100.

81. Microsoft Corporation, Weston S. foreach: Provides foreach looping construct. R package version 1.5.2.; 2022.

82. Weston S. Microsoft corporation. doParallel: foreach parallel adaptor for the ‘parallel’package. R package version 1.0. 17. 2022.

