## Supplementary material for "Population Size Mediates the Effects of Mating Systems on Evolutionary Rescue Under Worsening Environments" (from another version of this preprint: version 1, 2026-04-01)

**Mating Systems and Evolutionary Rescue**

**Table S1**: State Variables and Parameters

| Parameter/Variable | Description | Type | Values/Range |
| --- | --- | --- | --- |
| Population Variables |  |  |  |
| b | Probability of birth per offspring attempt | Fixed | 0.5 |
| K | Carrying capacity of the population | Fixed | 100, 250, 500 |
| t | Total number of time steps in the simulation | Fixed | 200 |
| e | Initial environmental value | Variable | 0.2 |
| mating_system | Type of mating system | Fixed | "monogamy" or "polygyny" |
| parental_care | Determines if offspring survival depends on mismatch and heterozygosity of both parents’ (TRUE) or only mother’s (FALSE) | Fixed | TRUE (for mutual mate choice monogamy) or FALSE |
| Individual Variables |  |  |  |
| O | Maximum number of offspring per female | Fixed | 5 |
| I | Age of sexual maturity | Fixed | 2 |
| grp.m | Number of males assessed by females per mating group | Fixed | 10 |
| grp.f | Number of females assessed by males per mating group | Fixed | 10 |
| alpha.m (α_m)_ | Strength of condition dependence for male sexual trait | Fixed | 2 |
| alpha.f (α*f*) | Strength of condition dependence for female sexual trait | Fixed | 2 |
| cost.m | Cost of sexual trait expression in males affecting survival | Fixed | 0 to 0.5 |
| cost.f | Cost of sexual trait expression in females affecting survival | Fixed | 0 to 0.25 |
| beta.m (*β_m_*) | Male mate preference strength | Fixed | 0 to 2.5 |
| beta.f (*β_f_*) | Female mate preference strength | Fixed | 0 to 5 |
| homozygosity penalty | Scaling factor penalizing low heterozygosity in survival, reproduction and signal trait | Fixed | 0, 0.25, 0.5, 0.75, 1 |
| mismatch.penalty | Scaling factor penalizing high mismatch in survival, reproduction and signal trait | Fixed | 0, 5, 10 |
| alive | Individual survival status | Variable | 1 (alive) or 0 (dead) |
| sex | Individual sex | Fixed | "M" (male) or "F" (female) |
| age | Individual age | Variable | 0 to 10 |
| Heterozygosity (*H*) | Proportion of loci where genetic strands differ | Variable | 0 to 1 |
| phenotype | Individual quantitative phenotypic trait value determining adaptation | Variable | Normal distribution, mean = 0.2 (initial) |
| Environmental Variables |  |  |  |
| e.var | Mode of environmental variation | Fixed | “Directional" |
| directional.rate | Mean change per time step | Fixed | 0 (stable environment), 0.0005, 0.001, 0.0015, 0.002, 0.0025, 0.003, 0.0035, 0.004, 0.0045, 0.005 |
| Genetic Variables |  |  |  |
| loci | Number of genetic loci for heterozygosity calculation in each strand | Fixed | 50 (default) |
| mutation_phe | Standard deviation of phenotypic mutation | Fixed | 0.01 (default) |
| mutation | Probability of allelic mutation (currently disabled in code) | Fixed | 0 (default) |

**Table S2**: Submodels and Mathematical Formulations

| Submodel | Description | Mathematical Formulation |
| --- | --- | --- |
| Environmental Variation | Updates the environmental value $e$based on the mode of variation. | For e.var = "Directional":  - If directional.rate = 0, $e=e_{0}+sample(\{-1,1\})\cdot U(0,0.02)$  Else if $i<50$, $e=e_{0}+sample(\{-1,1\})\cdot U(0,0.02)$  otherwise, $e=e+N(directional.rate,0.003)$ |
| Mismatch (*M*) | Computes mismatch as the absolute difference between phenotype and environment. | *\|e – phenotype\|* |
| Signal Trait | Calculates sex-specific signal trait for mature individuals ($age\geq I$). | For $age\geq I$:  Males: $\frac{0.5}{1+(mismatch.penalty\cdot M+inbreeding.penalty\cdot(1-H))\cdot\alpha_{m}}$-  Females: For $age<I$: $signal.trait=0$ |
| Death Probability | Determines survival probability based on density, mismatch, heterozygosity, and signal trait costs. | $D=density-\left( Mismatch penalty\times M+Homozygosity penalty\times\left( 1-H \right)+\left\vert signal trait \right\vert\times cost \right)$  density = *K* / population size, mismatch penalty and inbreeding penalty are variable, and cost (cost.m for males, cost.f for females,) represents the survival cost of expressing the signal trait. |
| Mate Choice | Assigns mating pairs based on signal traits, preference strengths, and mating system. | - Group assignment:  Females to $\lceil length(is.female)/grp.f\rceil$ groups;  males to $\lceil length(is.male)/grp.m\rceil$ groups.  Female choice probability: If $\beta_{f}>0$ and group size $n_{m}>0$,  $P_{i}=\frac{p_{i}^{-\beta_{f}}}{\sum_{j=1}^{n} p_{j}^{-\beta_{f}}},\quad for \beta_{f}>0$, where $p_{i}$is male rank; else $P_{i}=1/n_{m}$.  Male choice probability: if $\beta_{m}>0 group size n_{f}>0,$  $P_{i}=\frac{p_{i}^{-\beta_{m}}}{\sum_{j=1}^{n} p_{j}^{-\beta_{m}}},\quad for \beta_{m}>0$, where $p_{i}$is female rank; else $P_{i}=1/n_{f}$. |
| Reproduction | Determines number of offspring based on parental mismatch and heterozygosity. | Offspring penalty:  If parental_care = TRUE:  $offspring.penalty=Mismatch penalty\cdot\frac{M_{f}+M_{m}}{2}+Homozygosity penalty\cdot(1-\frac{H_{f}+H_{m}}{2})$  If parental_care = FALSE:  $offspring.penalty=Mismatch.penalty\cdot M_{f}+Homozygosity.penalty\cdot(1-H_{f})$  Number of offspring:  $offspring=\lfloor O\cdot(1-\frac{offspring.penalty}{offspring.penalty+1})\rfloor$ |
| Effective Population Size | Calculates effective population size based on variance in reproductive success. | Female effective size:  $N_{f_{e}}=\frac{1}{\sum(female\_offspring/total\_offspring)^{2}}$  Male effective size: $N_{m_{e}}=\frac{1}{\sum(male\_offspring/total\_offspring)^{2}}$  Effective population size: $N_{e}=\frac{4\cdot N_{m_{e}}\cdot N_{f_{e}}}{N_{m_{e}}+N_{f_{e}}}$if $N_{m_{e}}>0$ and $N_{f_{e}}>0$, else $0$. |

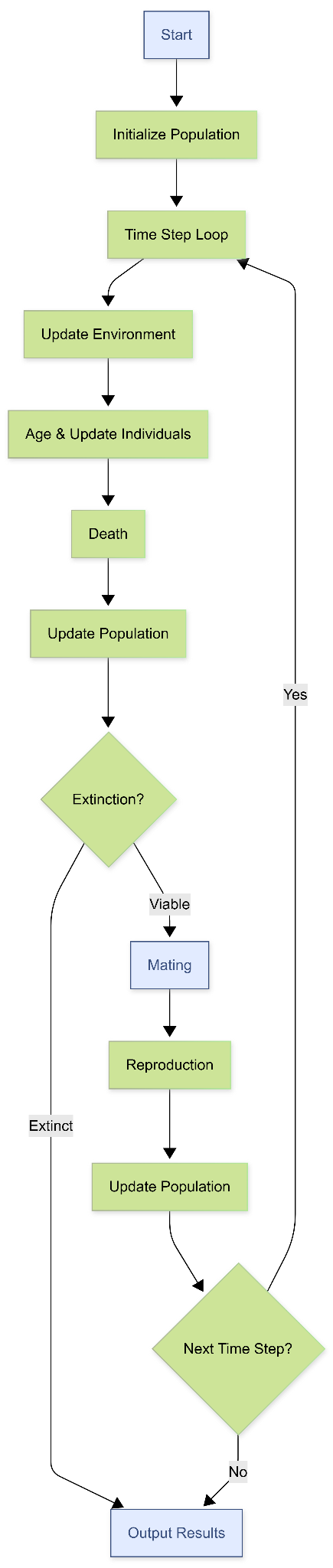

**Figure S1:** Flowchart explaining model work flow with major events in blue. Diagram created using [Mermaid.js](https://mermaid.js.org/).

A)
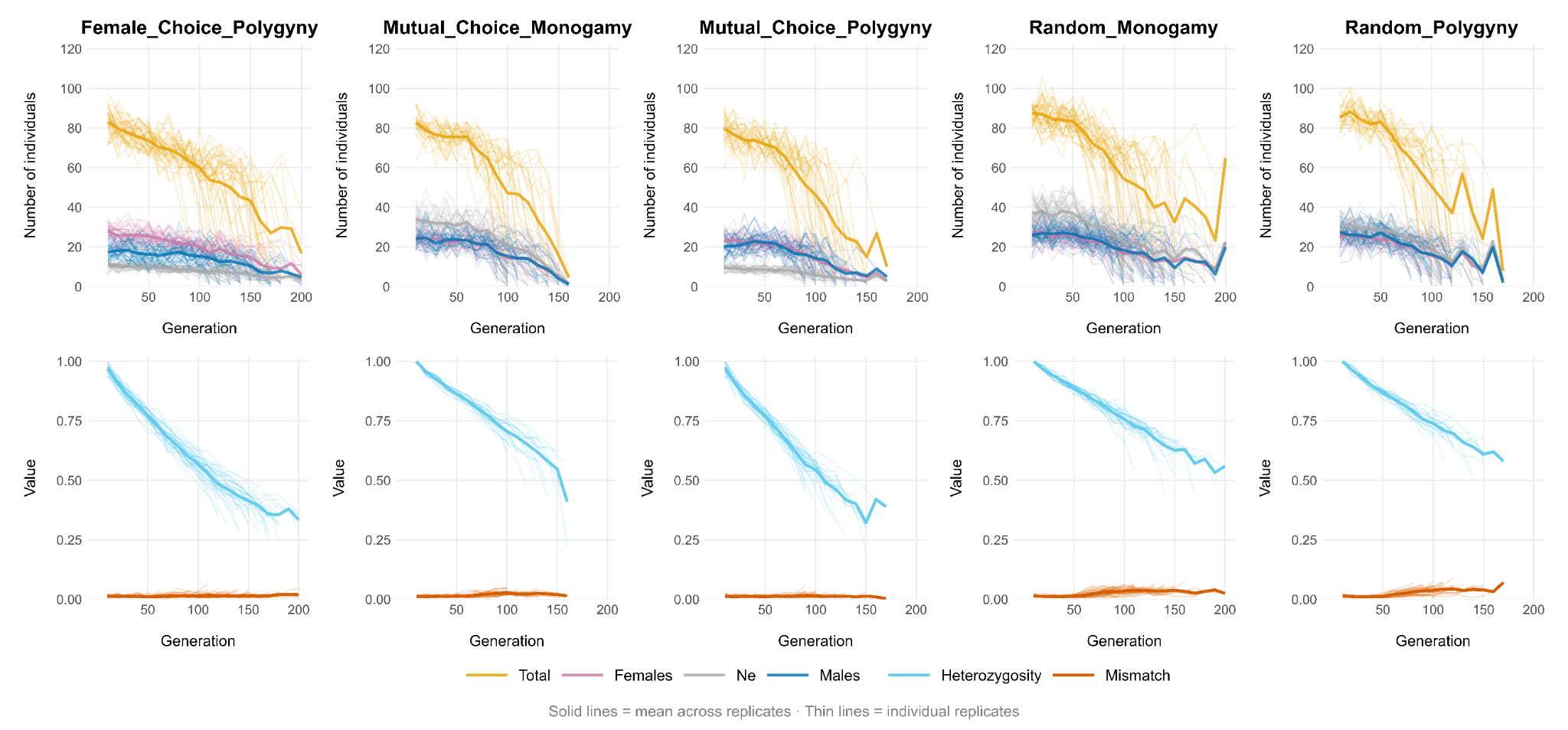

B)
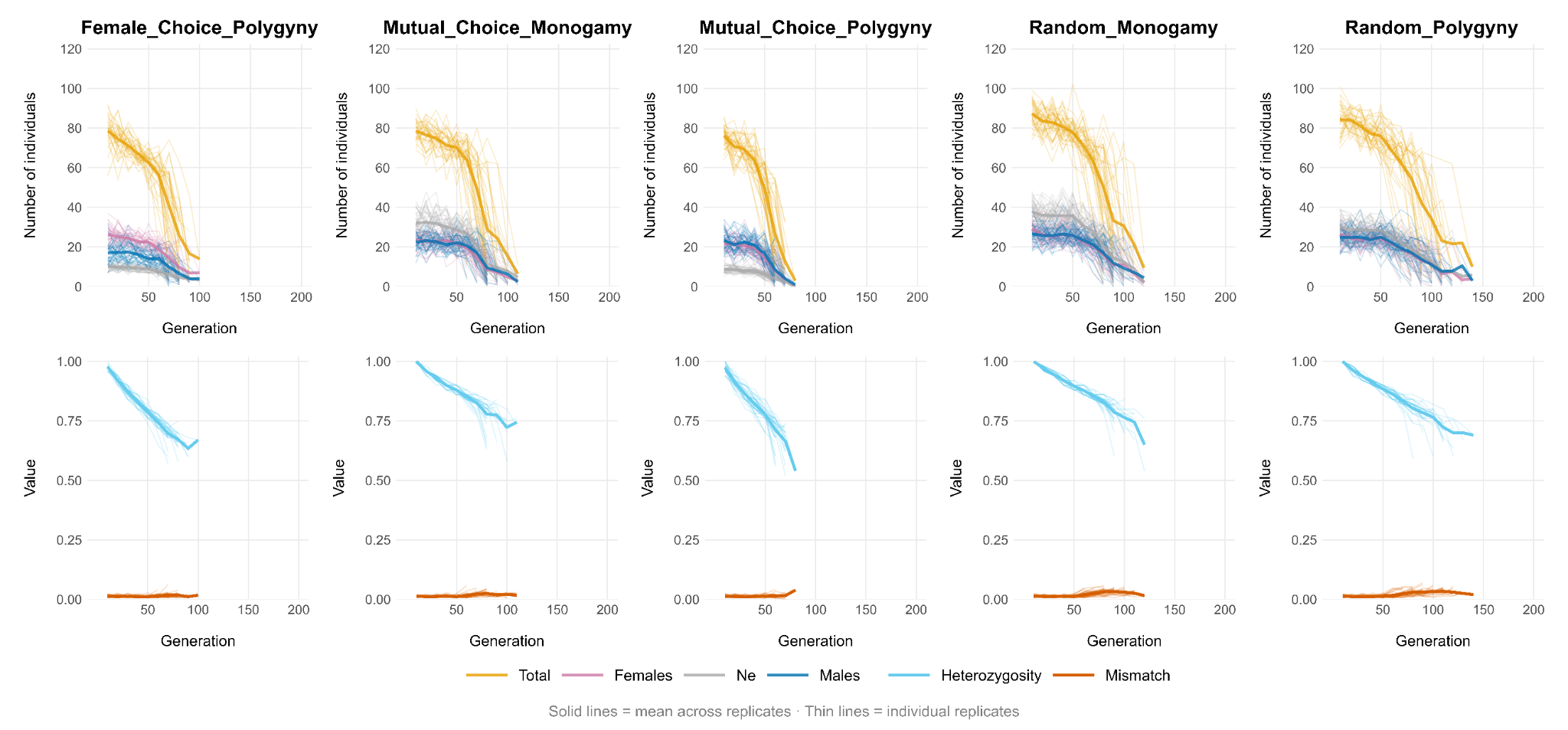

**Figure S2:** Demographic changes under carrying capacity *K* = 100, mismatch penalty = 5, directional environmental change rate = 0.002, and heterozygosity penalties of (A) 0.5 and (B) 1. Top panels in both (A) and (B) show mean (thick lines) and output from each simulation run (thin lines) population size dynamics plotted every ten time steps, including numbers of males (dark blue), females (pink), total population size (orange), and effective population size (gray). Bottom panels display the corresponding mean (thick lines) and output from each simulation run (thin lines) of median population heterozygosity (*H,* light blue) and environmental mismatch (*M,* red), also plotted every ten time steps. Each line represents the mean of 50 replicate simulations for all mating systems.

A)
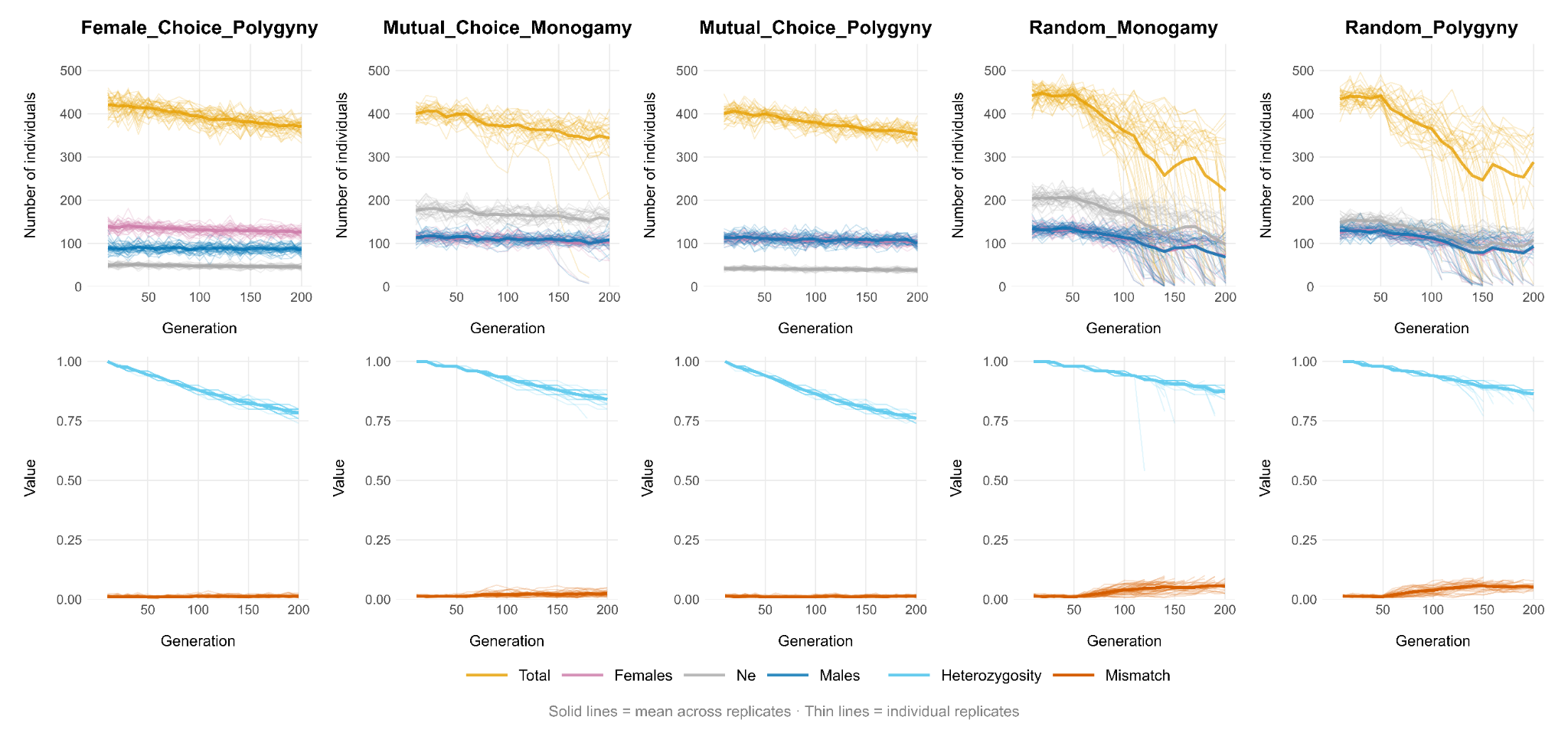

B)
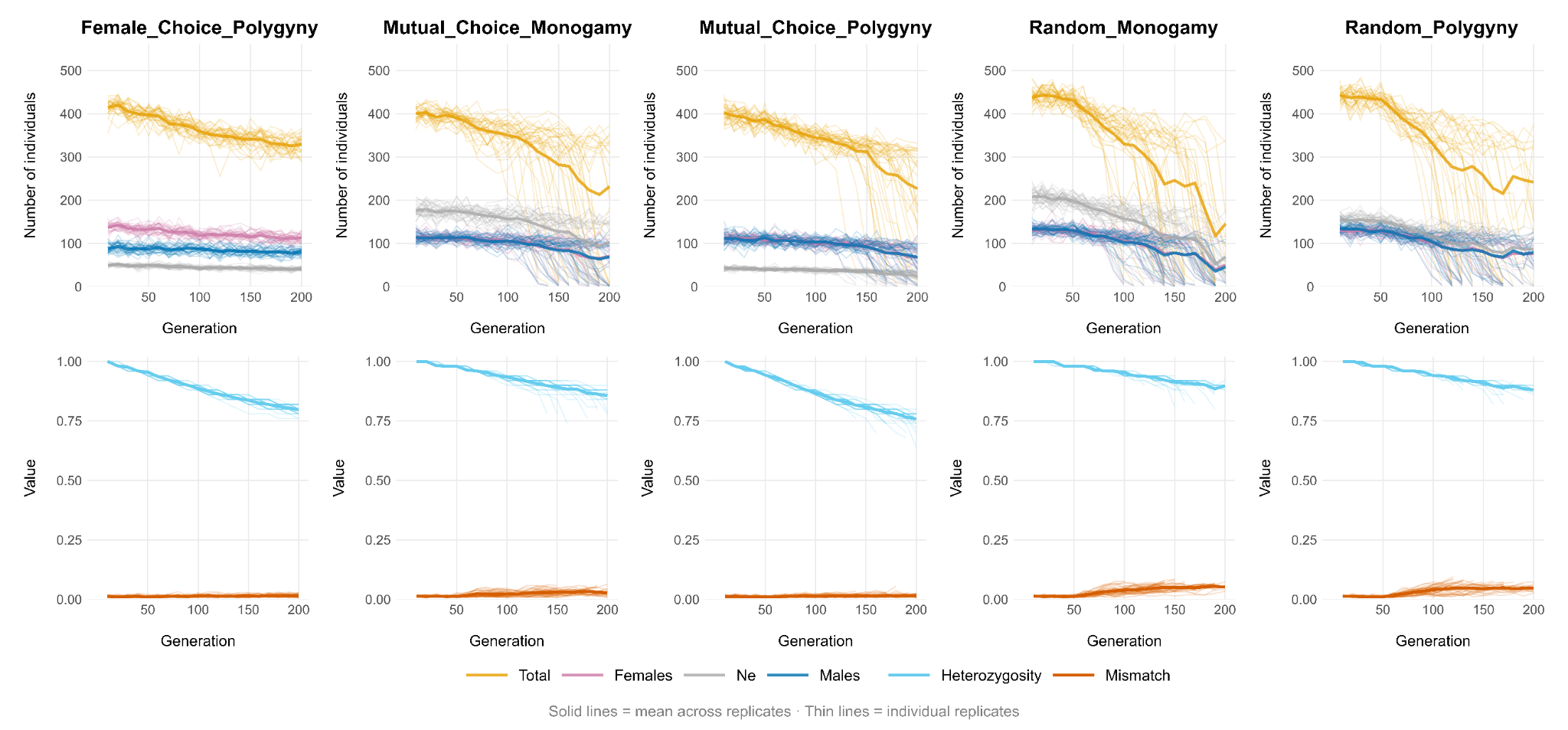

**Figure S3:**  Demographic changes under carrying capacity *K* = 500, mismatch penalty = 5, directional environmental change rate = 0.002, and heterozygosity penalties of (A) 0.5 and (B) 1.

Top panels in both (A) and (B) show mean (thick lines) and output from each simulation run (thin lines) population size dynamics plotted every ten time steps, including numbers of males, females, total population size, and effective population size. Bottom panels display the corresponding mean (thick lines) and output from each simulation run (thin lines) of median population heterozygosity (*H*) and environmental mismatch (*M*), also plotted every ten time steps. Each line represents the mean of 50 replicate simulations for all mating systems

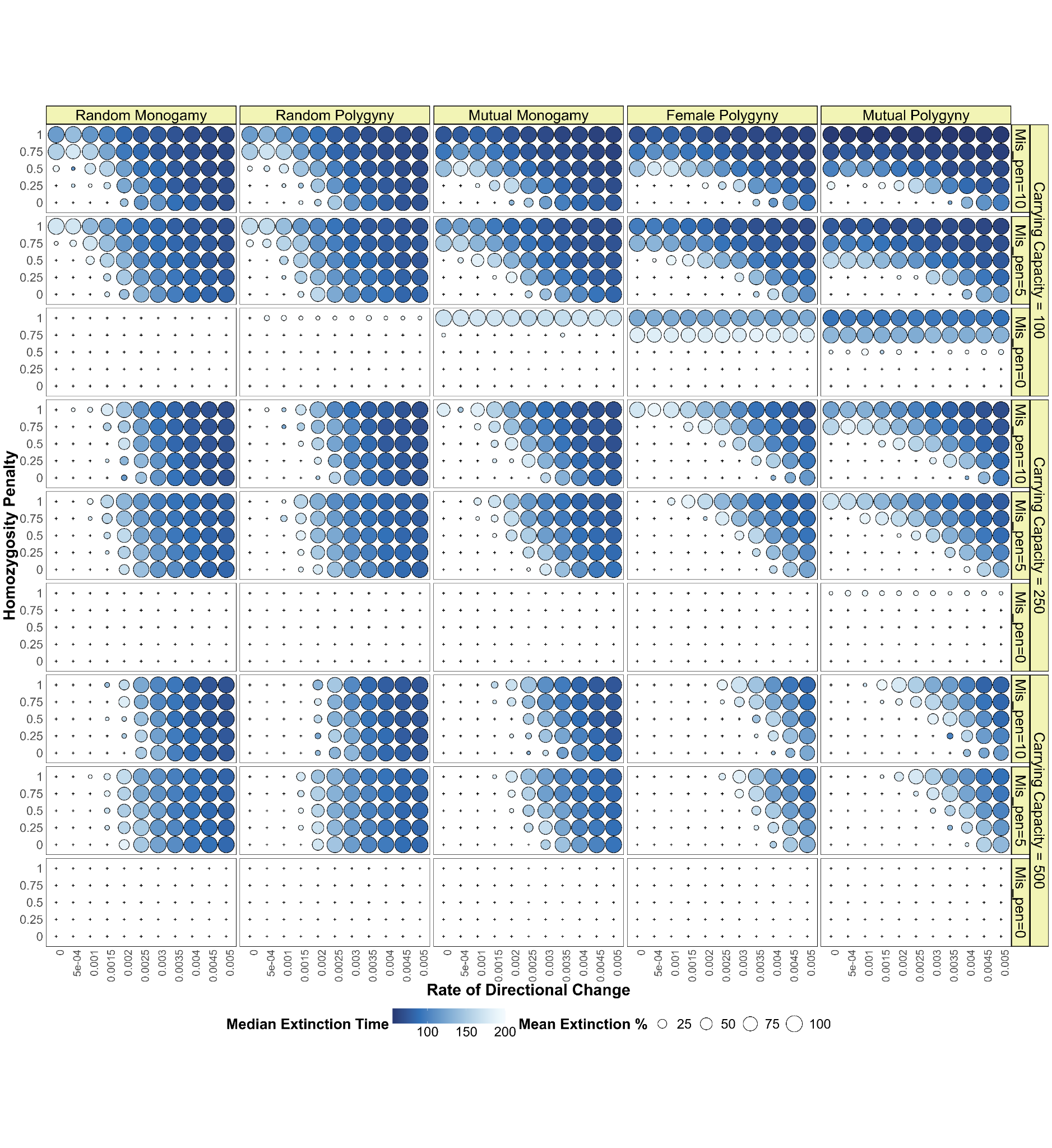

**Figure S4.** Heatmaps showing the percentage of populations that went extinct (size of the circle) and median timesteps when the populations went extinct (colour of the circle) across varying carrying capacities (*K*). Panels show results for *K* = 100, 250, and 500 from top to bottom. The x-axis represents rates of directional environmental change, and the y-axis represents inbreeding penalties. Vertical panels correspond to different mating systems, while horizontal panels vary in mismatch penalties (Mis_pen). The blocks with extinction probabilities of zero are denoted by “+”. Extinction probabilities were calculated from 50 simulations of 200 timesteps each for every combination of parameter values with parameters for mating systems described in Table 1.
